# Massive single-cell RNA-seq analysis and imputation via deep learning

**DOI:** 10.1101/315556

**Authors:** Yue Deng, Feng Bao, Qionghai Dai, Lani F. Wu, Steven J. Altschuler

## Abstract

Recent advances in large-scale single cell RNA-seq enable fine-grained characterization of phenotypically distinct cellular states within heterogeneous tissues. We present scScope, a scalable deep-learning based approach that can accurately and rapidly identify cell-type composition from millions of noisy single-cell gene-expression profiles.

---

Single-cell RNA-seq (scRNA-seq) can provide high-resolution dissection of complex biological systems, including identification of rare cell subpopulations in heterogeneous tissues, elucidation of cell-developmental programs, and characterization of cellular responses to perturbations^1–5^. Recent platforms, such as DropSeq^6, 7^, Microwell-seq^8^ and GemCode^9^, have enabled large-scale scRNA-seq on millions of cells at a time, which offers an unprecedented resolution at which to dissect cell-type compositions.

However, these advances have led to two acute challenges. First, single-cell profiles are highly susceptible to transcript amplification noise and dropout events^10, 11^, and such artifacts can become even more pronounced as tradeoffs are made to sequence larger numbers of cells. Second, current analytical packages^10–16^ are unable to scale to large datasets, including all cells and measured genes, due to computational memory and/or speed restrictions. New approaches are needed to extract informative representations from these extremely noisy, massive, high-dimensional scRNA profiles.

To overcome these challenges, we developed scScope, a software package that uses deep learning to extract informative features (low-dimensional representations of gene expression profiles) per cell from massive single-cell data (**Fig. 1a**). A major innovation of scScope is the design of a self-correcting layer. This layer exploits a recurrent network structure to iteratively perform imputations on zero-valued entries of input scRNA-seq data (**Methods**). In one joint framework, scScope conducts batch effect removal, cellular-feature learning, dropout imputation and parallelized training (when multiple GPUs are available) (**Supplementary Fig. 1**). To our knowledge, scScope is the first parallelized deep-learning framework for unsupervised, single-cell data modeling that can deal with both massive and noisy single-cell expression data.

**Fig. 1.**
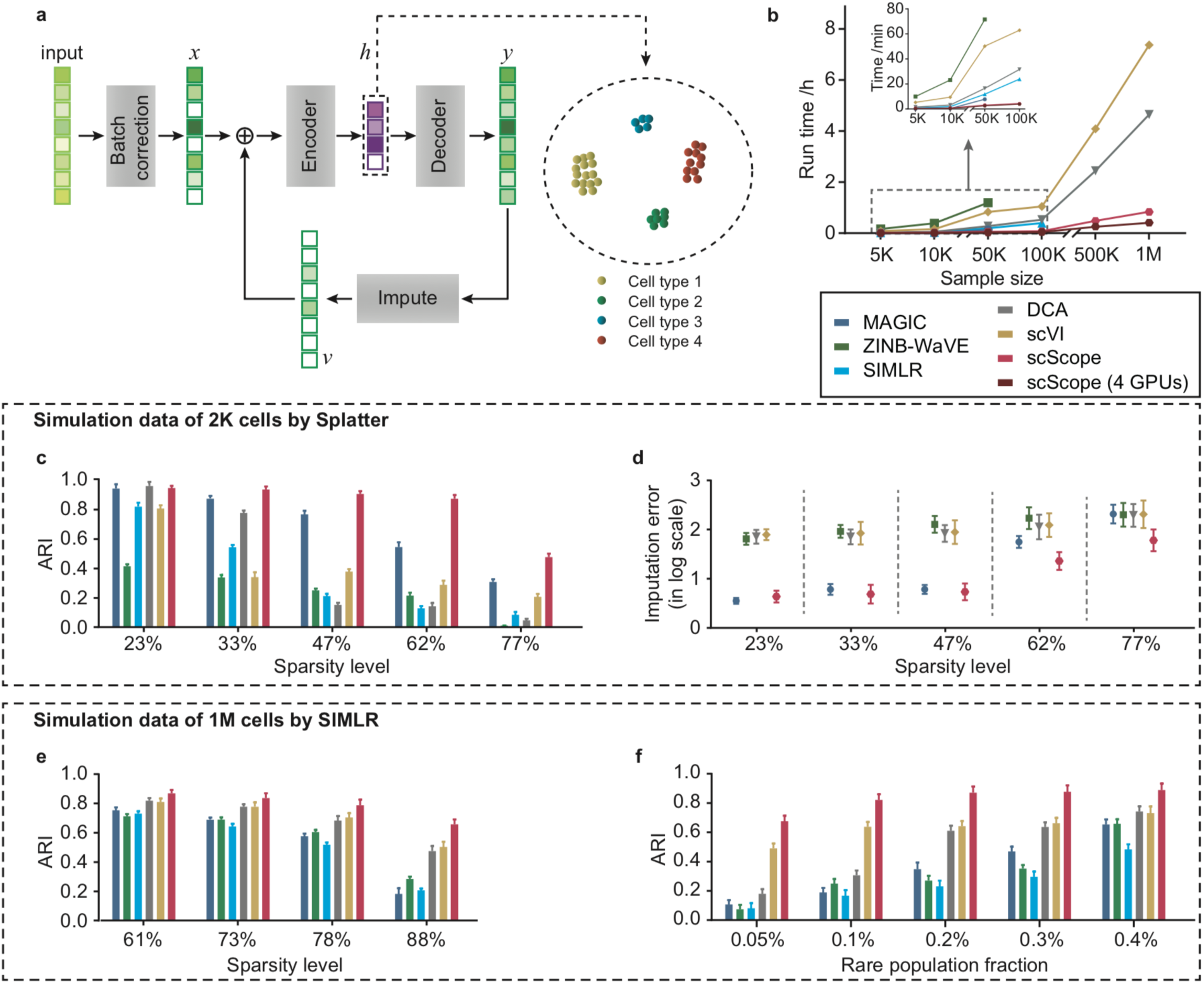
**Overview of scScope architecture and performance on simulated datasets.** **a) Overview of the recurrent network architecture of scScope.** An input single-cell profile with dropout gene measurements (white entries) is corrected for batch effects, then the corrected vector *x* is sequentially processed by an encoder layer (for feature extraction), decoder layer (for noise reduction) and imputation layer (for dropout imputation). The imputed vector *v* is added back to the batch-corrected input profile *x* to fill in missing values. This process proceeds recursively *T* times to produce a final signature feature vector output ℎ used for biological discovery, such as identification of phenotypically distinct subpopulations. **b) Comparison of run time on dataset of different scales.** Datasets of varying size were randomly subsampled from a dataset containing 1.3 million mouse brain cells and used for comparison (**Methods**). A training epoch was set to 100 for all deep learning methods (DCA, scVI and scScope), and a latent dimension of 50-dimension was used for all methods. **c**-**d**) **Comparison of clustering and imputation accuracy on 2K single-cell datasets generated by Splatter.** Methods were compared for varying fractions of sparsity levels (controlled by dropout rate, **Supplementary Table 2**). Accuracy measurement is based on (**c**) adjusted Rand index or (**d**) fractional measure of imputation errors. For each simulated condition, 10 random replicates were simulated; mean values and standard deviations (error bars) are reported. **e-f**) **Comparison of clustering accuracy on 1M single-cell datasets generated by simulation strategy as used in SIMLR.** Methods were compared for varying (**f**) fractions of dropout genes or (**g**) rare subpopulation fractions (10 replicates were used; ARI reported as in **c-d**). scScope and scVI directly analyzed the whole dataset, while other methods analyzed 20K down-sampled subsets.

We evaluated scScope on its scalability, ability to identify cell subpopulations, impute dropout gene information and correct batch effects. We used a variety of datasets in these evaluations, including two approaches for simulating data with “ground truth” and six biological datasets with varying degrees of size, complexity and prior biological annotation. Further, we compared the performance of scScope with three published “non-deep” methods (MAGIC^16^, ZINB-WaVE^11^ and SIMILR^12^) and two unpublished “deep” learning models (scVI^17^ and DCA^18^) (**Methods** and **Supplementary Table 1**). We note that among all compared methods, only scScope offers the ability for parallelized GPU training; for fair comparison we used a single GPU for comparisons unless noted otherwise (**Methods**).

We first tested the scalability and training speed of scScope on a mouse brain dataset^19^, which contained 1.3M cells, and we focused on the 1,000 most variable genes (**Fig 1b, Methods**). We evaluated the computational costs of all methods over a wide range of subsampled data sizes (from 5K to 1M; **Methods**). scScope was able to complete its analysis of the full dataset in under 50 minutes using a single GPU (this runtime can be significantly dropped by using multiple GPU training). In comparison, the non-deep approaches were unable to scale beyond 100K cells, and the deep approaches, while able to scale to 1M cells, required at least seven times more computing time than scScope. Thus, scScope is both scalable and highly efficient in dealing with large datasets.

To calibrate the accuracy of scScope on simulated datasets, we made use of two third-party packages for generating scRNA-seq data (**Methods** and **Supplementary Table 2**). First, we used Splatter^20^ to generate moderate-sized datasets of varying sparsity levels (percentage of 0-valued genes), containing: 2K scRNA-seq profiles with 500 genes, and three underlying subpopulations. In terms of discovering these underlying subpopulations, scScope performs similarly to other approaches when there are only minor dropout effects, but shows a large advantage in accuracy as dropout rates increase to realistic ranges observed in biological data^6, 8^ (**Fig. 1c**). In terms of imputation error, at low sparsity (<50%) scScope and MAGIC outperformed all other methods, though at high sparsity scScope outperformed all other approaches (**Fig. 1d**). A major contributor to the accuracy of scScope is its recurrent architecture (**Fig. 1a**), which allows imputed output to be iteratively improved through a selected number of recurrent steps (T). When T=1, the architecture reduces to a standard autoencoder^21^, and we found that T=2 provides a signification jump in performance. Overall, we found that T=2 offers the best tradeoff between speed and accuracy (**Supplementary Fig. 2**) and set this as the default parameter for our evaluations of scScope (though users have the option to change T for different applications).

Second, we used the simulation framework in SIMLR to generate massive-sized and more heterogeneous datasets of varying sparsity levels containing: 1M scRNA-seq profiles with 500 genes, and 50 underlying subpopulations. To perform our evaluations, the deep approaches were able to operate over the full datasets, while the non-deep approaches required down-sampled training strategies (**Methods**). We found that scScope consistently outperformed the other methods, particularly at high sparsity levels (**Fig. 1e**). An increasingly important task for scRNA-seq profiling approaches is to identify rare cell subpopulations within large-scale data. As might be expected, deep-learning approaches performed better than non-deep approaches, which required down-sampling. Overall, scScope consistently performed well on these unbalanced datasets, showing accuracy even down to subpopulations of size 0.05% of the overall dataset (**Fig. 1f**). Thus, our calibration showed that scScope can efficiently and accurately identify cell subpopulations from datasets with high dropout rates, large numbers of subpopulations, rare cell types and across a wide range of dataset sizes.

We next evaluated scScope on four experimental single-cell RNA datasets containing varying degrees of biological “ground truth”. These datasets were used to test the ability of scScope to: remove batch effects (**Fig. 2a**; lung tissue^8^), recover dropout genes (**Fig. 2b**; CBMC dataset^22^), identify minor subpopulations (**Fig. 2c-e**; retina dataset^6^), and test clustering accuracy for a large dataset with varying numbers of analyzed genes (**Fig. 2f**; mouse cell atlas^8^).

**Fig. 2.**
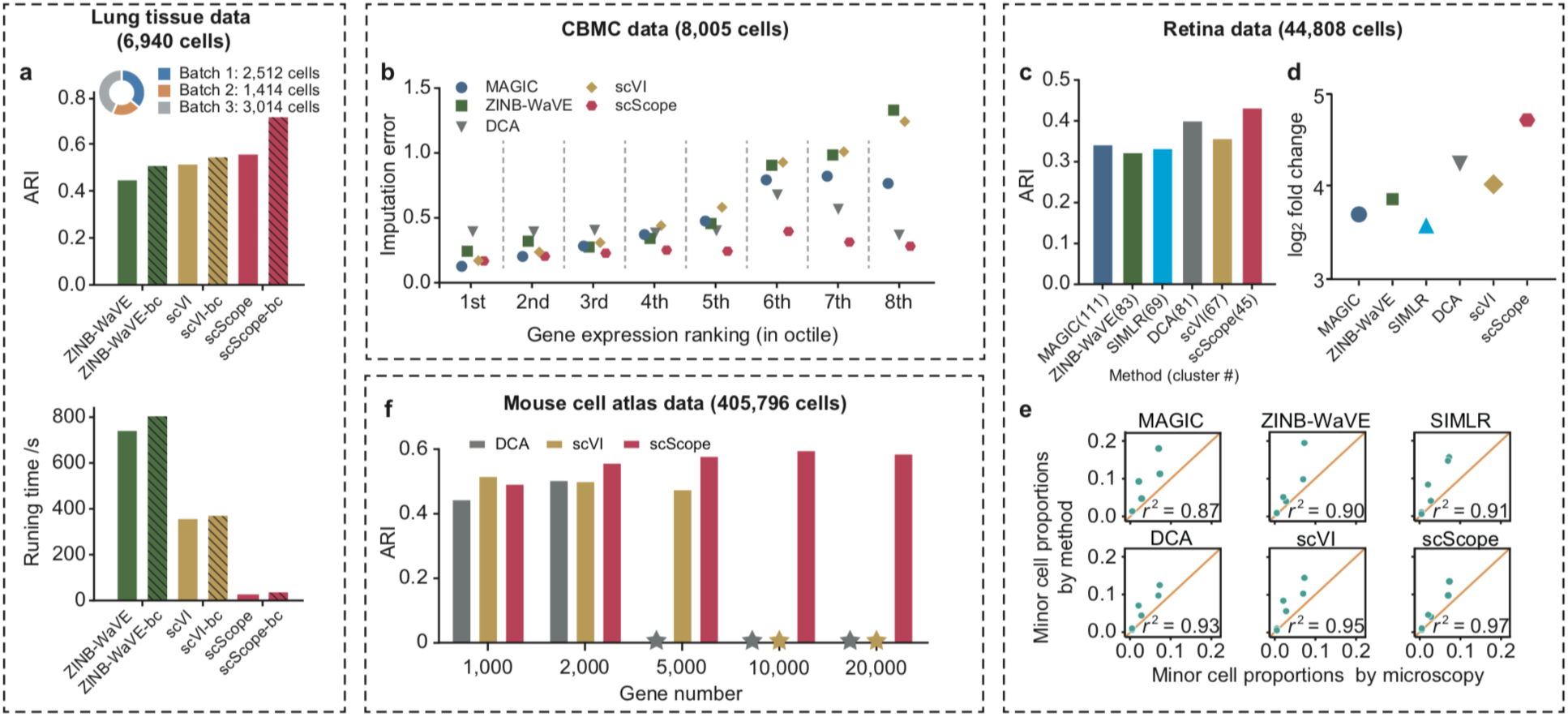
**Evaluation of methods on experimental scRNA-seq datasets.** **a) Analysis of batch correction.** Comparison on (top) clustering accuracy and (bottom) computational runtime without or with batch correction using mouse lung tissue scRNA-seq dataset. **b) Analysis of imputation accuracy for different gene expression levels.** Comparison of imputation error for dropout genes with different (octiles) gene expression level using the cord blood mononuclear cell (CBMC) scRNA-seq dataset. **c-e**) **Analysis of minor subpopulations discovery.** Using the retina scRNA-seq dataset, we compared: (**c**) the accuracy of cell-type identification (cluster numbers derived from each method are indicated in parentheses), (**d**) the significance of cell-type markers identified, and (**e**) the correlation of minor cell-type proportions identified by computation or microscopy (for all cell types, see **Supplemental Fig. 3**). **f) Analysis of subpopulation identification for increasing gene depth.** Using the mouse cell atlas, we compared the ability of different approaches to identify the 51 known tissues in the atlas. **\***: provided software package was unable to complete task.

To test the ability to remove batch effects, we made use of the lung tissue dataset (part of the mouse cell atlas), which contained ∼7K scRNA-seq profiles obtained from three different batches. We examined the 1,000 most variable genes and used PhenoGraph^23^ to identify subpopulations (**Methods**). Only ZINB-WaVE, scVI and scScope incorporated methodology for removing batch effects, and we tested their accuracies—with or without batch effect correction—for discovering 32 previously reported cell subpopulations. All three methods improved accuracy when the option for batch correction was enabled; scScope in particular showed a dramatic improvement in accuracy (**Fig. 2a**, top). Reassuringly, correcting for batch effects did not compromise the short runtime of scScope (**Fig. 2a**, bottom).

What is the dependency of imputation accuracy on gene expression level? We made use of the cord blood mononuclear cell (CBMC) dataset, which contained ∼8K scRNA-seq profiles and 1,000 most variable genes (**Methods**). Following the strategy used for scVI (**Methods**), we sequentially simulated dropouts for genes based on octile of expression ranking (**Fig 2b**). We found for imputing small count values that MAGIC and scVI performed well, while for large count values DCA worked well. However, scScope showed small imputation errors consistently across the entire range of expression.

Can minor subpopulations be identified? We made use of the mouse retina dataset^6^, which contained 44K cells from dissociated mouse retinas. We applied all methods to the 384 most variable genes and used PhenoGraph^23^ to identify subpopulations (**Methods**). The original study identified 39 cell subpopulations using fine-grained manual adjustment and expert visual inspection, which we took as a reference for our comparisons. Compared to other approaches, scScope automatically identified the most similar clustering (number and assignment) to those reported in the original study (**Fig. 2c**). We annotated the clusters to cell types based on gene markers reported in the original study (**Supplementary Table 3** and **Methods**). (The subpopulation of pericytes, identified by manual adjustment in the original study, was missed by all tested methods.) Overall, clusters identified by scScope showed the most statistically significant enrichment of specific cell-type markers (larger fold-changes) (**Fig. 2d**) and were highly consistent with previous, microscopy validation estimates of cell-type composition and proportion^24^ either without (**Fig. 2e**) or with (**Supplemental Fig. 3**) including the major cell type of rod cells. Thus, scScope performed well at recovering cell-type compositions even when cell proportions are unbalanced.

What is the benefit of analyzing increasing numbers of genes? We made use of the mouse cell atlas, which contained 400K cells sampled from 51 tissues. Only the deep learning algorithms were able to scale to these data sizes. To perform automatic identification of subpopulations on large datasets, we designed a scalable clustering approach (**Methods** and **Supplemental Fig. 4**). We made use of the 51 known tissue types to assess accuracy of the clustering results. Here, we found that all three algorithms performed similarly for 1,000 and 2,000 genes (**Fig. 2f**). However, the best performance overall was achieved by scScope by analyzing 10,000 genes. We note that for this analysis, we made use of an option in scScope’s software for scalable memory allocation (**Methods**). Thus, scScope offers the opportunity to analyze increased numbers of genes.

Finally, we applied scScope to investigate novel biology in datasets. We focused on the ability to reveal changes in cell-type composition under perturbed conditions (**Fig. 3a-c**; intestinal dataset^25^), and the ability to scale to large datasets and reveal new subpopulations (**Fig. 3d-e**; brain dataset^19^).

**Fig. 3.**
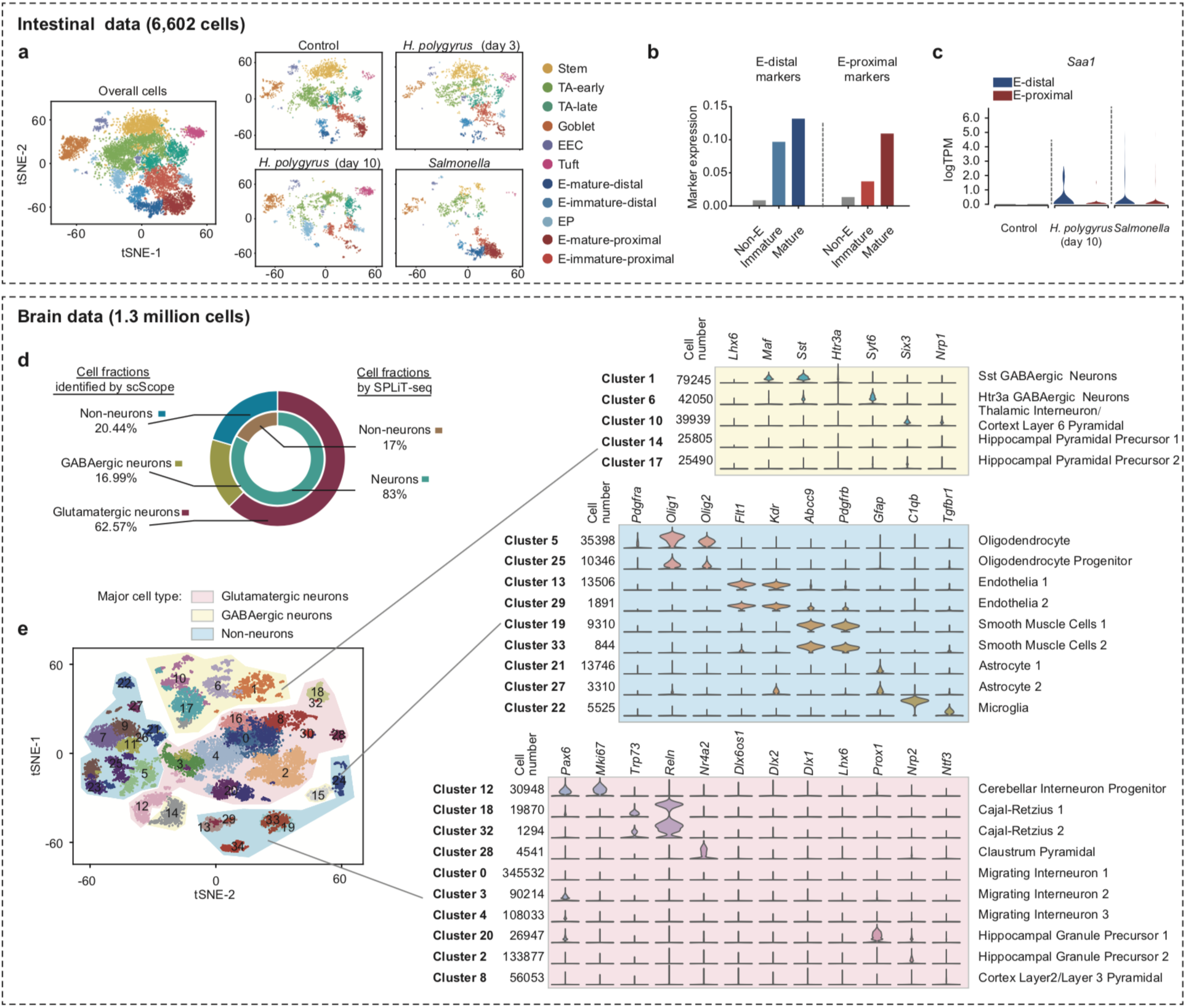
**Application of scScope to explore biology in two experimental datasets.** **a-c**) **Analysis of intestinal scRNA-seq dataset.** (**a**) Changes in cell-type composition of mouse intestinal epithelial under different infection conditions, visualized via tSNE plots. TA: transit amplifying, EEC: enteroendocrine, EP: enterocyte progenitor, E: enterocyte. scScope identified four subtypes of enterocyte cells. (**b**) Identification of mature vs. immature and distal vs. proximal enterocyte subpopulations. Shown are expression levels of E-distal and E-proximal gene markers (average UMI count) on the four enterocyte subtypes predicted by scScope and, for comparison, all other clusters (Non-E). (**c**) Discovery of differential expression of the gene *Saa1* in distal and proximal enterocytes after *Salmonella* and *H. polygyrus* infections. **d-e**) **Analysis of brain scRNA-seq dataset.** (**d**) Fractions of three major cell types (Glutamatergic neurons, GABAergic neurons and non-neurons) identified by scScope and comparisons with reported neuron fractions by previous SPLiT-seq research. (**e,** left) scScope results visualized using tSNE on 30K cells (randomly sampled from the full dataset). Clusters were divided to three major types based on gene markers. (**e,** right) Large-scale annotation of clusters to known cell types according to top 10 overexpressed genes. Vertical axis (left): clusters with known cell type annotations and corresponding cell numbers. Horizontal axis: differentially expressed marker genes across shown clusters. Vertical axis (right): previously reported cell subtype-specific genes, used to provide cluster annotation.

The intestinal dataset contained ∼10K cells obtained from mouse intestines with different infection conditions. We applied scScope to the 1,000 most variable genes (**Fig. 3a** and **Methods**). In the original study, enterocytes were identified as a single cluster. Interestingly, scScope further subdivided this cell type into four subpopulations: differential expression of markers provided delineation of distal *vs*. proximal enterocyte subpopulations, while expression levels of these markers provided further delineation into immature *vs*. mature subtypes. The assessment of maturity was based on expression levels of distal or proximal markers (**Fig. 3b** and **Supplementary Fig. 5**). Identifying these refined enterocyte subpopulations allowed us to make predictions about specific cell-type response to infection. For example, the pro-inflammatory gene *Saa1* was overexpressed during both *Salmonella* and *H. polygyrus* (day 10) infections in distal enterocytes, but not in proximal enterocytes (**Fig. 3c** and **Supplementary Table 4**). This geographic pattern of *Saa1* expression is known for *Salmonella* infection, but is a novel prediction arising from scScope analysis for *H. polygyrus* infection. Thus, scScope can be used to rapidly explore scRNA-seq data, predict novel gene function and identify new cell subtypes from perturbed conditions.

The 1.3M cells in the brain dataset were obtained from multiple brain regions, including the cortex, hippocampus and ventricular zones, of two embryonic mice. scScope automatically identified 36 clusters, and we assigned each cluster to one of three major cell types based on criteria from the Allen Brain Atlas (http://brain-map.org) (**Fig. 3d-e**, **Supplementary Table 5** and **Methods**): Glutamatergic neurons, GABAergic neurons and non-neuronal cells. The proportions of neurons and non-neurons identified by scScope were consistent with cell proportions reported by previous brain study^26^ (**Fig. 3d**). We investigated whether we could identify biological meaning to the 36 clusters, some of which contained fewer than 1,000 cells. Satisfyingly, by comparing our top overexpressed genes with known cell-type markers^26–28^ (**Supplementary Table 6**), we were able to assign two thirds of the clusters to known cell types (**Fig. 3e**). Thus scScope can rapidly, automatically and directly identify *bona fide*, rare cell types from datasets with over a million single-cell transcriptional profiles.

Taken together, scScope offers a platform that will keep pace with the rapid advances in scRNA-seq, enabling rapid exploration and accurate dissection of heterogeneous biological states within extremely large datasets of single-cell transcriptional profiles containing dropout measurements.

## Acknowledgements

We thank Drs. Jeremy Chang, Satwik Rajaram and Laura Sanman for their helpful comments. We gratefully acknowledge the support of NIH R01 EY028205 and GM112690, NSF PHY-1545915 and SU2C/MSKCC 2015-003 to SJA, NCI-NIH RO1 CA185404 & CA184984 to LFW and the Institute of Computational Health Sciences (ICHS) at UCSF to SJA and LFW.

## Author contributions

Y.D., F.B. and Q.D. developed the deep learning algorithms. Y.D. and F.B. conducted experimental analysis on both simulated and biological datasets. The manuscript was written by Y.D., F.B., L.F.W. and S.J.A. All authors read and approved the manuscript.

## Methods

### 1. scScope model and training

#### Architecture

The architecture of the scScope network has four modules (**Fig. 1a**). The parameters in these layers are learned from data in an end-to-end manner through optimization. We note that scScope is flexible in terms of normalizing and scaling of input data, as long as the input values are non-negative.

##### Batch Effect Correction

scSope offers the option to correct for batch effects, following the same batch effect correction mechanisms used in ref 11. We denote: the input single-cell profile as 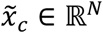; the number of batches by *K*; a one-hot experimental batch effects indicator vector 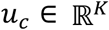(the one-hot (i.e. non-zero) entry indicates the corresponding batch from which 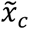 was sequenced); and the learnable batch correction matrix as 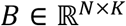. Then, we remove the batch effect from 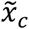 via:
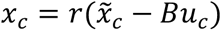
Throughout, we use *r*(∙) to denote the standard rectified linear unit (ReLU):
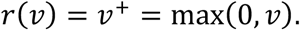
This rectifier enforces the property that *x_c_* only has non-negative values, which is expected for actual gene count data.

Then, *x_c_* is regarded as the single-cell profile with batch effects removed. We note that *u_c_* is an optional input for end-users to indicate each cell’s batch identity. In scScope, *u_c_* is set by default to a zero vector, assuming there is no batch effect among cells.

##### Encoder

For each cell *c*, scScope makes use of an encoder layer to compress the high-dimensional single-cell expression profile 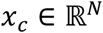 into a low-dimensional, latent representation 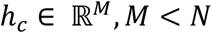. Here,
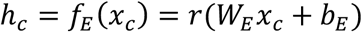
where the encoder layer *f_E_*(∙) is a composition of a linear transform with learnable parameters 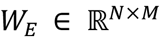 and 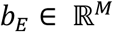 followed by the nonlinear activation *r*(∙). We note that ReLU is the most widely used non-linear function in deep learning due to its ease in back-propagating gradient information across layers.

##### Decoder

A decoder layer is established to decompress the latent representation *h_c_* to an output 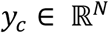 that is of the same dimension as the input single-cell profile,
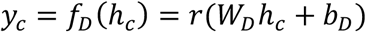
where the decoder layer *f_D_*(∙) is a composition of a linear transform with learnable parameters 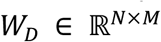 and 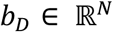 followed by the nonlinear activation *r*(∙). The nonlinear ReLU activation *r*(∙) sets all negative values to zero, which makes experimental sense for gene transcript abundance measured by nonnegative values. When minimizing the differences between *y_c_* and 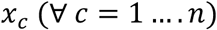 in the training set (of size *n*), the above encoder-decoder network acts as a typical auto-encoder (AE) neural network. AEs exhibit the potential for generating a “clean” signal *y_c_* by removing additive noise from *x_c_* via *L_p_* norm minimization (typically, *p* = 2)^29^. However, the AE paradigm lacks a mechanism for dealing with dropout entries (missing data), which can be an even more severe problem than noise for scRNA-seq.

##### Imputer

We next focus on estimating missing data measurements. Zero-valued measurements can be due either to biological reasons (un-transcribed genes), or to technical reasons (dropouts). Our imputation step is designed to deal with the later problem, namely to estimate dropout gene expression values. Of course, a challenge shared by all single-cell-based imputation is to distinguish between zero-expressed *vs.* dropout measurements.

To this end, we developed a self-correcting layer to impute missing entries caused by dropout during sequencing. Our imputation approach implicitly makes use of the fact that subsets of genes are often co-regulated (*e.g.* by common transcription factors or pathway activation) and that their patterns of co-expression can be learned by observation of sufficiently many cells. The decoder in our AE framework is enabled to find such patterns from the latent space representation. The studies of transcriptional regulatory mechanisms have led to the development of many approaches for network-based pathway recovery.

For our purposes of imputing expression values from initial zero values, we chose to make use of a widely used sparse graph-reconstruction approach^30^. We only perform imputation operations for zero measurements of the original single cell profile *x_c_*. Assume for cell *c* that gene *g* has zero expression value in the expression profile *x_c_*. Our goal is to impute *v_cg_*, the value of gene *g* in cell *c*, using the output of the decoder layer, *y_c_*. We do this by:
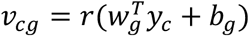
where learnable parameters 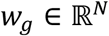 is a sparse vector and *b_g_* is a bias for each gene *g*. (The *i^th^* entry in vector *W_g_* indicates the influence of a gene *i* on the expression of gene *g*.) These gene-specific parameters are learned from regression across all cells. We used *y_c_* rather than the original input *x_c_* to represent gene expression values as it is expected to contain less expression noise after AE reconstruction. As above, the nonlinear ReLU activation *r*(∙) forces regressed values to be non-negative; the hope for truly zero-expressed genes is that regression will return non-positive values, which are set to zero after ReLU activation.

As with our previous network layers, we can describe the imputation step in matrix form as follows. We define *Z_c_* (*resp*. 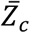) as the set of zero-(*resp*. non-zero-) valued genes in profile *x_c_* of the *c*-th cell. We can now write the imputation layer as:
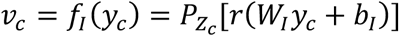
where the imputation layer *f*_1_(∙) is the triple composition of a linear transform (with sparse weight matrix 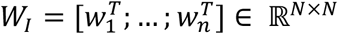 and bias 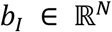), an ReLu *r*(∙), and “entry-sampling” operator 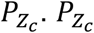 sets to zero all entries not in *Z_c_*, as we are only interested in imputation for zero-valued genes in the profile *x_c_* (*i.e.*, for any vector *r*, 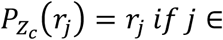 *x_c_* and 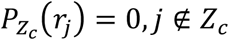).

After obtaining imputed vector *v_c_*, we obtain a corrected single-cell expression profile: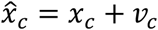. This corrected single cell profile 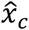 can be used as the input to the encoder layer to learn an updated latent representation through the encoder-decoder framework. Because missing values have been estimated in 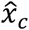, the new latent representation is expected to be better than the original one encoded from 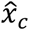.

#### Learning the objective function

The imputation layer imposes a recurrent structure on the scScope network architecture. For clarity of exposition, the recurrent scScope can be unfolded into multiple time steps (**Supplementary Fig. 6** shows three steps). Then, the whole recurrent scScope framework can be described as:
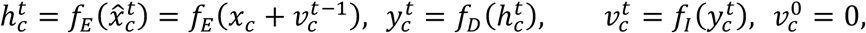
for iterations *t* = 1 … *T*. At the first step, the correcting layer’s output 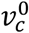 is set as zero. We note that if we only consider one learning step (*T* =1) then scScope is a standard normal auto-encoder network.

The learning objective for scScope is defined by the unsupervised, self-reconstruction pursuit (as typically used in auto-encoder training):
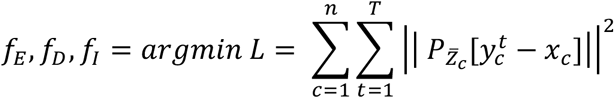
The entry-sampling operator 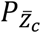 forces loss computation only on (non-zero) measured entries of *x_c_*. The parameters in the encoder layer (*f_E_*), decoder layer (*f_D_*) and imputation layer (*f*_1_) are all learned by minimizing the above loss function. We note that scScope learns its informative latent representation 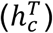 in an unsupervised manner. scScope requires no extra information (*e.g*. cell type) to accomplish its deep-learning training.

#### Multiple GPU training

scScope offers the option to train its deep model by using multiple GPUs, which can dramatically reduce runtime (**Supplementary Fig. 1**). In this mode, scScope replicates its network structure on multiple GPUs and aggregates all network parameters on the CPU. These network parameters include the connections and biases of all encoder, decoder and imputation layers of scScope. In a round of batch training, one GPU grabs the current network parameters from the CPU to use for its own network replicate of scScope. Then, for gradient calculation, the GPU processes a randomly chosen batch of *m* (= 64 or 512) single-cell expression profiles from a total of *n* single-cell profiles.

We apply a conventional gradient calculation framework for neural networks, which iteratively performs feed-forward and back propagation steps. In the feed-forward (FF) step, a GPU passes its batch of *m* single-cell samples through its locally stored scScope network and accumulates the losses for this batch. In the back propagation (BP) step, batch-dependent gradient information for network parameters on different layers is calculated by sequentially propagating accumulated loss from the end to the first network layer. This BP operation is performed by using gradient calculation functions wrapped in deep-learning packages (in our case TensorFlow). We apply this process independently across all *k* GPUs in a parallelized manner to obtain gradient information from a total *k* × *m* samples. The gradient information of those *k* GPUs is averaged by the CPU, *i.e*. 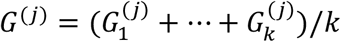, where 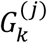 is the gradient calculated from the *k^th^* GPU in the *j^th^* round of optimization iteration. Finally, we apply adaptive moment estimation (ADAM)^31^ with default TensorFlow^32^ parameters to update the network parameters stored on the CPU. Iterations were terminated when either the objective function showed little changed (*i.e.* < 0.1%) or the number of iterations reached a maximal epoch (*e.g.* 100).

#### Cell subpopulation discovery

Representations outputted by scScope can be integrated with any clustering methods. Here we used the graph-based method PhenoGraph^23^, as it performs automated robust discovery of subpopulations, as well as determines subpopulation numbers automatically.

##### 1) Graph clustering for moderate-scale data

We directly applied the PhenoGraph software to datasets of moderate scale. All clustering results were obtained using a Python-implemented PhenoGraph package (version 1.5.2). We followed the suggested setting and considered 30 nearest neighbors when constructing graphs.

##### 2) Scalable graph clustering for large-scale data

scScope enables the feature learning on millions of cells. However, PhenoGraph is unable to handle millions of cells due to the extreme computational costs (**Supplementary Fig. 7**) and memory requirements in graph construction. To leverage the power of graph clustering on analyzing these large-scale data, we designed a density down-sampling clustering strategy by combining *k*-means and PhenoGraph.

In detail, we divided cells into *M* groups with equal size and performed *K*-means clustering on each group independently (**Supplementary Fig. 4**). The whole dataset was split to *M* × *K* clusters and we only input the cluster centroids into PhenoGraph for graph clustering. Finally, each cell was assigned to graph clusters according to the cluster labels of its nearest centroids.

In our implementation on the dataset of 1.3 million mouse brain cells, we took *M* = 500 and *K* = 400, which made it possible to process millions of data in only tens of minutes without loss of accuracy (**Supplementary Fig. 8** and **Methods)**.

##### 3) Scalable memory allocation for analyzing large numbers of genes in large datasets

For large datasets and gene numbers, scScope implements a scalable memory allocation strategy that allows the dataset to be broken into a smaller number of batches that can be loaded directly into memory. We note that when this option is used, minibatches are only selected from within each batch during training. This option was only used in **Fig. 2f** for the case of ≥10K genes; here the 400K mouse cell atlas dataset was broken into four batches of size 100K.

### 2. Imputing dropout genes

This output layer of the scScope neural network provides a complete set of gene profiles with all entries filled. Accordingly, we directly used filled values of this layer to impute missing results.

### 3. Implementation of comparison methods

We note that a hyper-parameter across all methods is the number of latent feature dimensions M (**Methods**); we observed that all methods were reasonably robust to changes in M (e.g. **Supplementary Fig. 9**), and to avoid an intractable number of possible comparisons, we set M=50 for all comparisons. Unless otherwise noted, the software packages were used as follows.

*Markov Affinity-based Graph Imputation of Cells (MAGIC):* The MAGIC algorithm was performed using the python-based package *magic*. We input the raw data and employed the `library_size_normalize()` function provided by the software for all simulated and real data to learn 50-dimension latent features.

*Zero-Inflated Negative Binomial-based Wanted Variation Extraction (ZINB-WaVE)*: We employed the R package zinbwave, and all default parameters were used to learn the 50-dimension feature vector. For running without batch correction, we set batch information for each cell as the same, and for batch correction mode the batch indices were input as reference.

*Single-cell interpretation via multi-kernel learning (SIMLR)*: We used the Python implementation of SIMLR with the authors’ default parameter settings. SIMLR needs to take the desired cluster number as input. For our simulated dataset, we input the known cluster numbers. For the retina dataset, where the “true” cluster number is unknown, we set it to 39, which is the cluster number reported in the original study^6^.

*Deep count autoencoder (DCA)*: DCA is an unpublished software based on TensorFlow framework. We installed the Python package of DCA (download date: Sep. 4, 2018) and ran DCA by setting the latent dimension to 50. We set the training epoch to 100 and kept all other default parameters.

*Single-cell Variational Inference (scVI):* scVI is an unpublished work on biorxiv. We used the Torch-based Python package of scVI (download date: June 5, 2018). In the original demonstration of the software, they assigned different parameters for different test datasets. Here, we set the training step to 0.001, epoch to 100 and the latent dimension to 50. The software shuffled the cell order randomly in training and did not offer the option to output the latent representation with the same input cell orders. In order to keep track of the input cells for later analysis, we appended cell IDs to the cell labels.

*scScope*: scScope was implemented in Python 3.6 with *TensorFlow-GPU* 1.4.1, *Numpy* 1.13.0, *Scikit-learn* 0.18.1 packages and was tested on a server with 4 GPUs (Nvida Titan X) and 64GB memory. For all experiments presented in this paper, we extracted 50-dimensional representations with 2 recurrent learning steps (*T* = 2). For medium-scale data size, we use the default batch size of 64 and epoch of 100 in training; for large-scale data set, we use a larger batch size of 512 and smaller epoch of 10 to speed up the learning process. Unless noted specifically, scScope was run on single GPU mode in our comparisons.

*Down-sampling training strategies on large-scale dataset:* It is not possible to run directly the non-deep-learning based approaches on dataset with millions of cells. For these packages, we randomly down-sampled datasets containing more than 1M cells to a subset of 20K cells. On these down-sampled datasets, single-cell feature vectors were learned by the respective method and clustered by PhenoGraph. A support vector machine (SVM) was trained on this subset in the latent feature space and then used to assign labels for the rest of cells in the unsampled dataset. The deep-learning approaches we tested can learn features on millions of cell profiles, but the software does not provide a function for automatic clustering on such large-scale datasets. Therefore, for comparisons we passed the output of their deep learning algorithms for single-cell feature learning to our scalable graph clustering approach for large-scale clustering.

All compared methods were run on the same server with Xeon E5 CPU, 64 GB memory, Nvidia Titan X GPU and Ubuntu 14.04 operation system. Further, all comparisons were performed using log transformed input (we observed similar relative performance of the six compared methods above across five different choices of input scaling or normalization methods; **Supplemental Fig. 10**).

### 4. Evaluation of clustering performance

We used the adjusted Rand index (ARI)^33, 34^ to compare label sets of two clustering methods. For two clustering results *U* and *V* with *r* and *s* clusters on a data set of *n* cells, *n_ij_* denotes the number of cells shared between cluster *i* in *U* and cluster *j* in *V*. And ARI is defined as
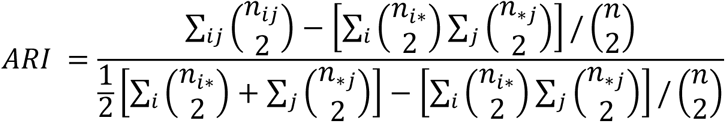
where 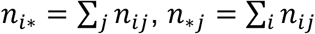 and *n* is the number of cells in the data set.

### 5. Evaluation of Imputation performance

The imputation accuracy was defined as the normalized distance between the imputed log count entries and log count ground truth entries. We constructed lists *l* and 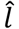 whose elements correspond to either ground truth or imputed values (respectively) for all dropouts entries across all cells (**Fig. 1d**). We defined the normalized error as:
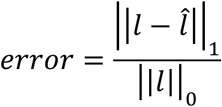
where 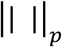 means the *l^p^* norm of a vector.

On the simulated data, the ground truth dropout vector *l* is known. However, for real biological data, the ground truth values for missing genes are unknown. To evaluate scScope’s imputation accuracy on real biological data, we followed the same down-sampling strategy as used for scVI^17^. Namely, we randomly split the entire collection of *n* cells into *n_train_* training cells and *n_val_* validation cells. We used the different imputation methods to build gene models from the *n_train_* cells. On each of the *n_val_* cells, we randomly set *p*% of its non-zero genes as “simulated” missing genes and set their corresponding count values to zero. The real measured values of these simulated missing genes were then used to generate the ground truth list *l*, and the list 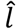 was based on inferred values for the simulated missing genes from the *n_val_* cells. The imputation error was calculated as above.

### 6. In silico studies

#### Simulation with Splatter

The simulation package Splatter^20^ is designed to generate realistic scRNA-seq data. We used this package to generate data with 2000 cells, 3 subpopulation groups, and dropout rates from 1 to 5.

#### Simulation with SIMLR

We used the simulation approach from SIMLR^12^ to generate large-scale scRNA-seq data due to limitations in scalability of Splatter. Following previous studies, we initially tested the performance of scScope for cell-subpopulation discovery using simulated data^10, 12^. We assumed the high-dimensional single-cell expression data 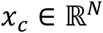 is generated or controlled by a latent code 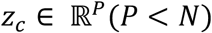, which is not observable. *Z_c_* is sampled from a Gaussian mixture model with *k* Gaussian components, *i.e.* 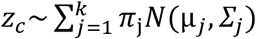. The mixture coefficients *π_j_* were chosen from a uniform distribution and normalized to sum to 1, the mean vector 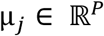 was uniformly sampled in [0,1]*^p^*, and the covariance matrix 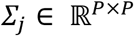 was chosen to be Σ = 0.1 × *I*, for identity matrix *I*.

To simulate single-cell gene vectors, we generated a projection matrix 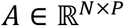 to map the low-dimensional latent code to a high-dimensional space. First, we simulated ground truth, 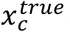, which cannot be observed:
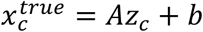
where each entry in *A* was independently sampled from the uniform distribution *U*(−0.5,0.5) and bias *b* = 0.5 is added to avoid negative gene expression in the high-dimensional mapping. Second, we simulated the observed profile, 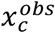, which may contain gene-specific noise and dropout artifacts due to the sequencing technique and platform. Noise was added to the true gene profile by: 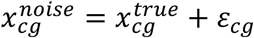, where 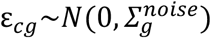 and 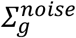 was uniformly sampled in the range of [0, 0.2] independently for each gene. Dropout events were added via a double exponential model with decay parameter^35^ *α*:
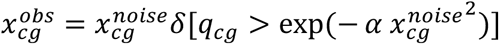
where *x_cg_* denotes the *g^th^* gene of *x_c_*, *q_cg_* was randomly sampled in [0, 1], and *δ* = 1 if its argument is true and = 0 otherwise. This double-exponential model is motivated by the widely-used assumption that low-expressed genes have higher probability to be influenced by dropout events.

We use the aforementioned generative model to create *N* single-cell profiles 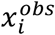, *i* = 1 … *N*. In the reported results (**Fig. 1e-f**), we repeat such random generation process for 10 times on *n* =10K and 1M simulated single-cell profiles under various conditions for performance comparisons among different approaches.

#### Simulation with rare cell subgroups

To generate cell subpopulations with rare cell types, *i.e.* cell clusters with very limited numbers of cells compared to the major clusters, we sample the mixture coefficients *π_j_* from a non-uniform distribution as
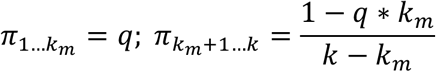
where 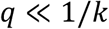 is the mixture fraction for each minor cluster, *k_m_* is the number of rare cell subpopulations, and *k* is the total number of subpopulations. With this minor modification from the previous section, we generate imbalanced cell-type compositions with *k_m_* rare cell types.

In our simulation results reported in **Fig. 1f**, we generate large-scale, imbalanced datasets with 1M single-cell profiles according to the following parameters: *α* = 0.5 (dropout rate), *k* = 50 (total number of clusters), *k_m_* = 5 (the number of rare cell types) and varied the rare cell-type mixture rate *q* in the range from 0.1%∼0.4%. 10 replicates for each condition were analyzed. We randomly down-sample these 1M single cell profiles to 10K. On the subsampled datasets, after learning features, we applied PhenoGraph for *de novo* cell subpopulation discovery. On the un-subsampled datasets, we used our scalable graph clustering approach (**Methods**). We note that subpopulation numbers are automatically determined by these clustering approaches.

### 7. Analysis of biological data sets

#### (a) Lung tissue data

The lung tissue dataset is part of the Mouse Cell Atlas data^8^. This dataset was downloaded from Gene Expression Omnibus (GEO) database (accession number: GSE108097). In this dataset, 6,940 cells were sequenced via three independent experiments by Microwell-seq, with 2,512, 1,414 and 3,014 cells in each batch. 1,000 high variable genes were selected for analysis. Then three batch correction methods (ZINB-WaVE, scVI and scScope) were used on the normalized data to learn the low-dimensional features. Cell types were identified by PhenoGraph clustering.

To evaluate the clustering accuracy, identified cell labels were compared with previously reported labels (https://satijalab.org/seurat/mca.html).

#### (b) Cord blood mononuclear cells dataset

In this dataset, 8,617 cord blood mononuclear cells (CBMC) were profiled by CITE-seq, a new technology which enabled the simultaneous measurement of protein levels and transcriptome levels for each cell^22^. This dataset was downloaded from GEO database (accession number: GSE100866). Cell types in CBMC have been extensively studied and identified. Based on this prior knowledge, 13 monoclonal antibodies were chosen to identify *bone fide* cell types. These antibodies serve as an “orthogonal” ground truth to evaluate analysis results based on RNA-seq data.

In the data pre-processing stage, we removed the spiked-in mouse cells and only kept 8,005 human cells for analysis using the cell-filtering strategy introduced in original study^22^. The top 1,000 most variable human genes were selected for downstream analysis after the log1p transformation and normalization by library size. For antibody data, we used the centered log ratio transformed antibody-derived tags (ADTs), which is also provided by authors.

To evaluate the performance of each method, we first automatically identified cell populations based on ADTs data using PhenoGraph. Then, scRNA-seq data were input to each method to learn latent representations which were used by PhenoGraph to predict cell types. The ADTs-derived cell types were taken as ground truth to evaluate the accuracy of cell types by scRNA-seq data.

#### (c) Retina data

In this dataset, 44,808 cells were captured from the retinas of 14-day-old mice and sequenced by DropSeq^6^. Data were obtained from the GEO database (accession number: GSE63473). In order to be comparable with published results, we followed previous experimental procedures to select 384-most variable genes^6^ and then to log transform their expression (log(TPM + 1)). After clustering, we annotated clusters obtained by each method using the same maker genes in the original study^6^.

We identified candidate cell types based on the highest average type-specific marker expression (**Supplementary Table 3**). For each cluster, we calculated the fold-change values of all cell-type markers, and if at least one of a type-specific gene marker was expressed significantly higher (log_2_ fold change > 0.5) than in all other clusters, we assigned the cluster with the candidate cell type. Otherwise the cluster was assigned to the cell type “Rod cell”.

#### (d) Mouse cell atlas data

The mouse cell atlas (MCA) dataset is designed to offer a comprehensive investigation of all major cell types in mouse^8^. Data were downloaded from the GEO database (accession number GSE108097). In the dataset, 405,796 cells were sampled from 51 tissues and were sequenced by Microwell-seq.

Data were firstly normalized by library size and 1,000, 2,000, 5,000, 10,000 and 20,000 top-variable genes were selected to test the scalability of each method on gene numbers. Only the deep-learning based methods (DCA, scVI and scScope) could be applied directly to this large-scale dataset. Further, to identify clusters in the MCA dataset, we applied our scalable clustering approach to the latent features.

In most of the 51 tissues, one major cell type dominated the cell population (see Figure 2b-c in ref 8). Therefore, we used the tissue identify as a proxy for ground truth to evaluate cell-type discovery.

#### (e) Intestinal data

In this dataset, intestinal epithelial cells were captured from 10 mice and sequenced using droplet-based scRNA-seq^25^. Data were downloaded from the GEO database (accession number GSE92332). Among all cells, 1,770 cells from 2 mice were infected by *Salmonella* for 2 days; 2,121 cells (2 mice) and 2,711 cells (2 mice) were infected by *H. polygyrus* for 3 and 10 days, respectively. An additional 3,240 cells were sequenced from 4 healthy mice as a control group. We again followed the same procedure that log-transformed the expression data and selected top 1,000 most variable genes as input to scScope. For cell subpopulation annotation, we first assigned clusters to one of 7 major cell types (stem, cell-cycle related, distal enterocyte, proximal enterocyte, goblet, enteroendocrine, and tuft) according to the maximum averaged expression of cell-type makers (**Supplementary Table 7**). Second, cell-cycle related clusters were subdivided into increasing stages of maturation (transit amplifying early stage, transit amplifying late stage, and enterocyte progenitor) based on the ratio of cell-cycle & stem cell markers to enterocyte expression (**Supplementary Table 8**). Third, the distal and proximal enterocyte clusters were further classified (immature *vs.* mature) based on increasing expression levels of the enterocyte gene markers.

After annotating clusters, we calculated the cell proportion for each mouse and then averaged the proportions among mice of the same infection condition. For significant tests of proportion changes after infection, we compared proportions of mice in control group and in infection group using a two-sided t-test and rank-sum test. P-values were obtained under the null hypothesis that no changes happened in proportions after infection.

Overexpressed genes for each cluster were also identified by the same differential expression analysis.

#### (f) Brain data

Data were obtained from 10x Genomics (http://10xgenomics.com). 1,308,421 cells from embryonic mice brains were sequenced by Cell Ranger 1.3.0 protocol. We transformed unique molecular identifier (UMI) count data into log(TPM+1)^6^ and calculated the dispersion measure (variance/mean) for each gene. According to the rank of the dispersion measure, we selected the top 1,000 most variable genes for analysis.

Due to the massive scale of the dataset, we set the batch size of scScope to 512 and trained the model for 10 epochs. Cells were further clustered into 36 groups by our density down-sampling clustering. We annotated clusters to three major types (excitatory neurons, inhibitory neurons and non-neuronal cells) based on maximal-expressed maker genes (**Supplementary Table 6**).

To identify cluster-specific overexpressed genes, we then conducted differential expression analysis for each gene. We normalized UMI-count to the range of [0 1] for each gene, enabling comparisons across genes. Then gene-expression fold-change and rank-sum P-values were calculated between cells within vs. outside each cluster. Significantly overexpressed genes were identified using the criteria of log_2_ fold change > 0.5 and rank-sum P-value < 0.05.

Data set was download from https://support.10xgenomics.com/single-cell-gene-expression/datasets/1.3.0/1M_neurons on December 10, 2017.

The data analysis by 10xgemonics was obtained from http://storage.pardot.com/172142/31729/LIT000015_Chromium_Million_Brain_Cells_Application_Note_Digital_RevA.pdf.

### 8. Evaluation of the density down-sampling accuracy

To evaluate the accuracy of the proposed density down-sampling clustering method, we randomly selected a subset (*n* = 5,000, 10,000, 20,000, 50,000, 100,000) of 1.3 million data and directly input corresponding scScope features to PhenoGraph for clustering. Then, we compared subset labels with labels obtained by density down-sampling clustering on the whole dataset. The comparison was repeated 100 times on different randomly selected subsets (**Supplementary Fig. 8**).

### 9. Software used in study

MAGIC: https://github.com/KrishnaswamyLab/MAGIC

ZINB-WaVE: https://github.com/drisso/zinbwave

SIMLR: https://github.com/bowang87/SIMLR_PY

DCA: https://github.com/theislab/dca

scVI: https://github.com/YosefLab/scVI

PhenoGraph: https://github.com/jacoblevine/PhenoGraph

### 10. Code availability

scScope can be obtained as an installable Python package, which can now be obtained via “pip install scscope”, and is available under the Apache license. All software, instructions and software updates are maintained on https://github.com/AltschulerWu-Lab/scScope.

