## Supplementary material for "Massive single-cell RNA-seq analysis and imputation via deep learning"

#### Supplementary Figures

**Supplementary Figure 1.** Training of scScope is implemented on multiple GPUs to enable fast learning and low memory cost (**Methods**). For each epoch, data batches are input into different GPUs for local parameter updates. Local parameters are summarized on the CPU for overall parameter updates and sent back to the GPUs for parameter learning in the next epoch. This architecture is enabled by TensorFlow functionality.

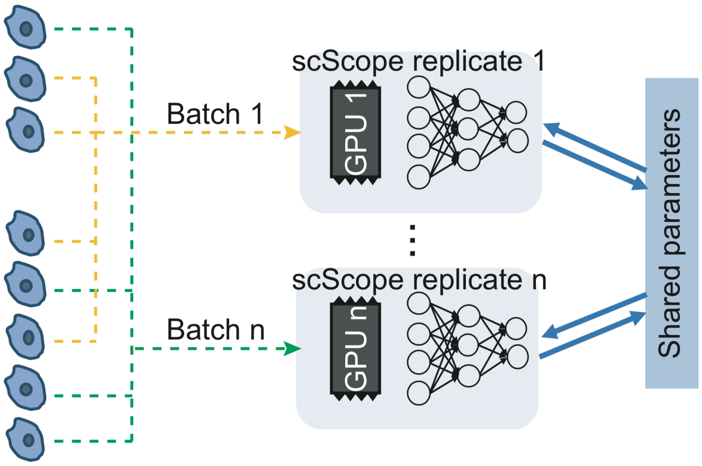

**Supplementary Figure 2.** Performance analysis of scScope under different recurrent iterations (T) using simulated data (top panel) and biological data (bottom). Clustering accuracy (left), imputation error (middle) and run time (right) were reported for each dataset.

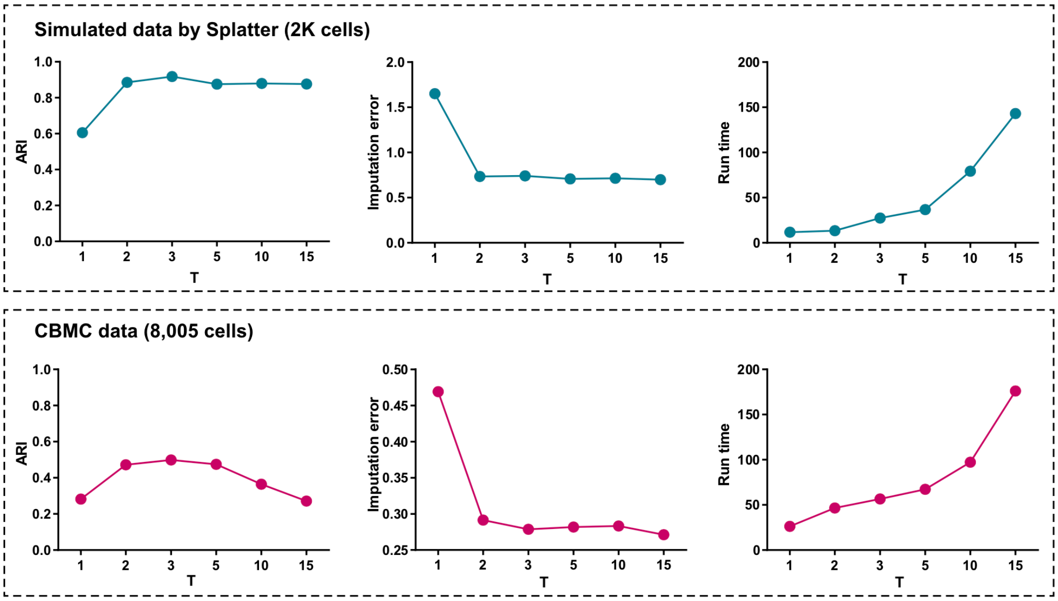

**Supplementary Figure 3.** The correlation of overall cell-type proportions identified by computation and microscopy on mouse retina dataset (as in **Fig. 2e**, but with all subpopulations).

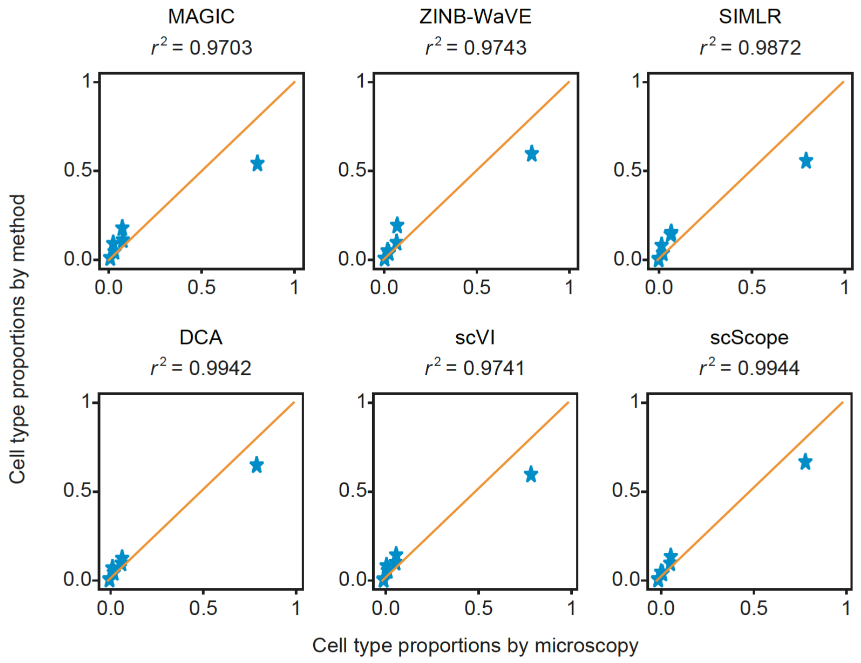

**Supplementary Figure 4.** Overview of scalable clustering approach for grouping large-scale data.

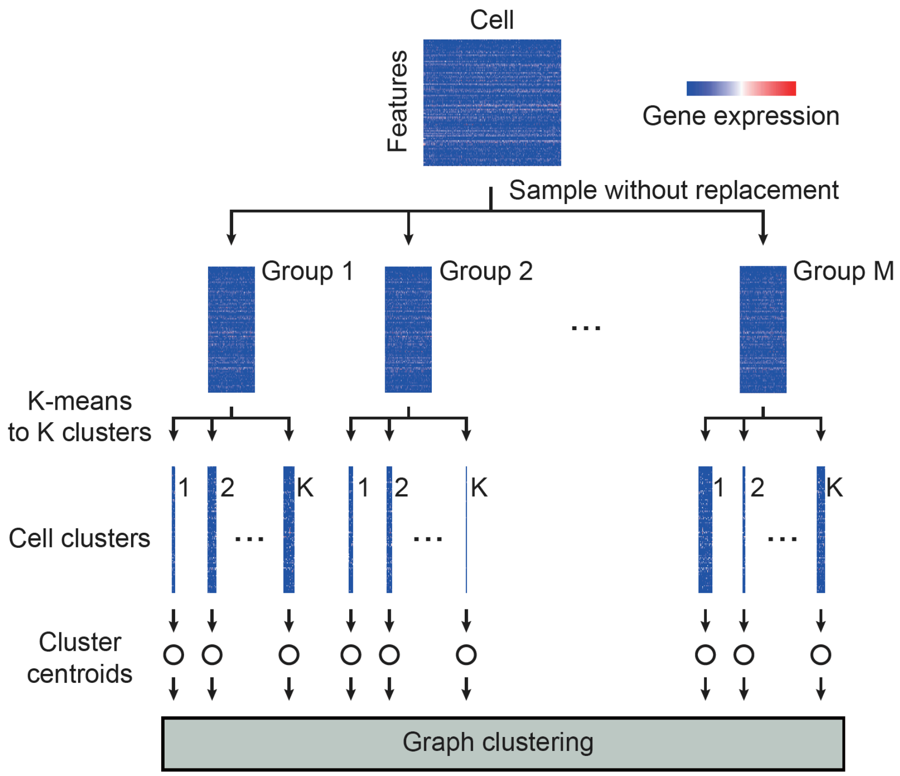

**Supplementary Figure 5.** Statistical evaluation (P-values) of differences between immature *vs.* mature, non-enterocyte (non-E) *vs.* immature, non-E *vs.* mature for E-distal and proximal clusters on mouse epithelial cell dataset.

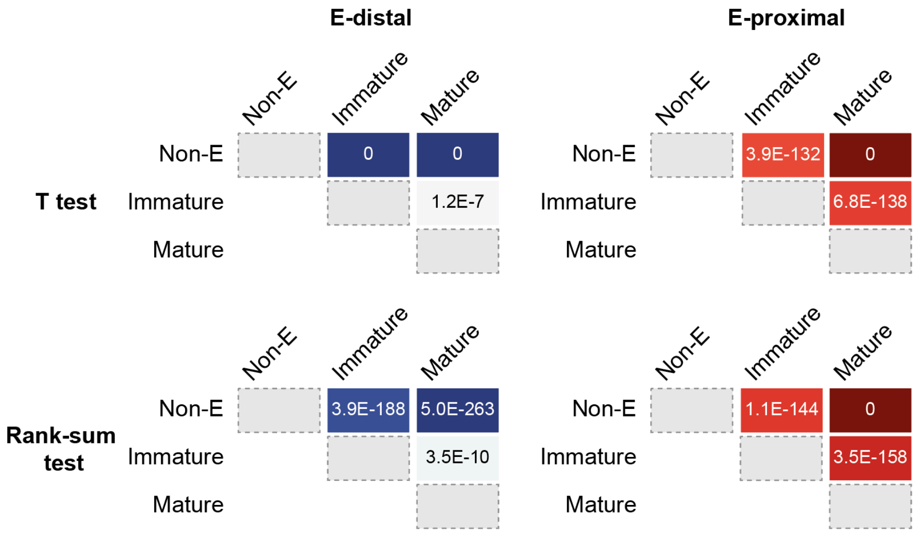

**Supplementary Figure 6.** scScope's recurrent network shown unfolded to three steps.

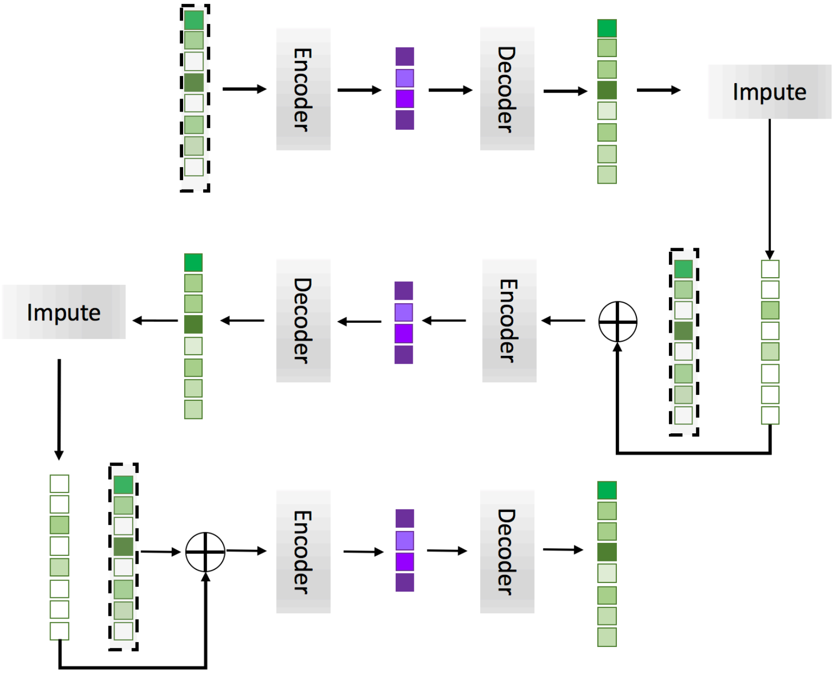

**Supplementary Figure 7.** Time costs of running PhenoGraph on simulated datasets of different scales. For data size in 5K ~ 50K, reported time costs were obtained as average values on 100 random repeats. For 100K or 200K data sizes, 10 repeats were used to report average time costs.

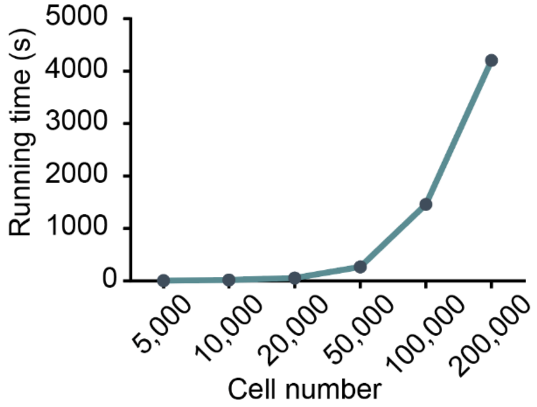

**Supplementary Figure 8.** Evaluation of the accuracy of our scalable graph clustering result. A subset of $n$ cells ($n$ = 5K, 10K, 20K, 50K and 100K) were randomly sampled from the 1.3 million mouse brain data set. The corresponding scScope features were clustered directly using PhenoGraph. Adjusted Rand index (ARI) were used to evaluate consistency between density down-sampling cluster labels and PhenoGraph labels. Mean ± STD of ARI were reported on 100 random repeats (only 10 repeats were used for $n$ = 100K).

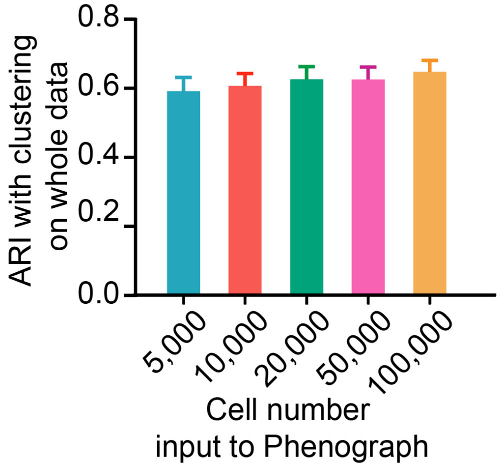

**Supplementary Figure 9.** Evaluation of performance for varying latent dimensions (M = 10, 30, 50, 100) using the 8K CBMC dataset.

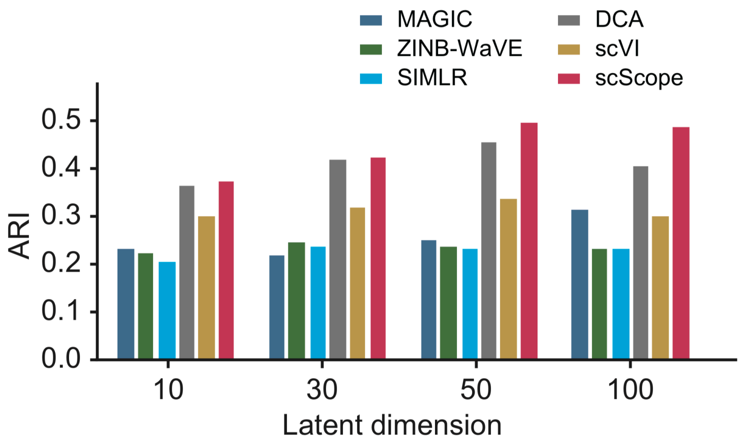

**Supplementary Figure 10.** Performance of comparing methods on different normalization/scaling methods using the CBMC dataset. The default normalization methods for MAGIC and DCA ("Library size") were disabled for the other four comparisons.

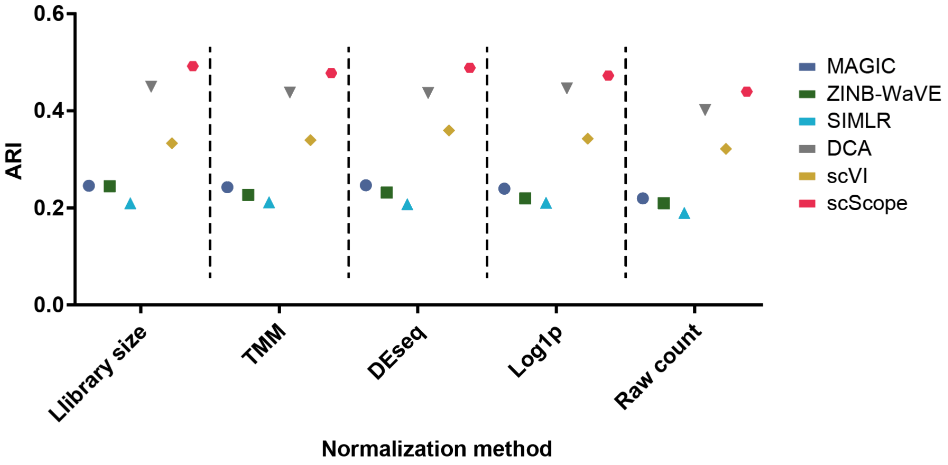

### Supplementary Tables

**Supplementary Table 1.** Summary of methods.

| | Underlying machine learning model | Dropout Imputation | Batch correction | Efficient large scale cell clustering | GPU-parallelized implementation |
|---|---|---|---|---|---|
| MAGIC | Linear algebra | Yes | No | No | No |
| ZINB-WaVE | Matrix factorization | Yes | Yes | No | No |
| SIMLR | Kernel learning | No | No | No | No |
| DCA | Deep neural network | Yes | No | Yes | No |
| scVI | Bayesian inference | Yes | Yes | Yes | No |
| scScope | Recurrent neural network | Yes | Yes | Yes | Yes |

**Supplementary Table 2.** Parameter settings and corresponding sparsity levels of simulated datasets by two scRNA-seq simulation tools (Splatter and SIMLR)

Splatter simulation

| Dropout rate | 1 | 2 | 3 | 4 |
|---|---|---|---|---|
| Sparsity level | 0.23 | 0.33 | 0.47 | 0.77 |

SIMLR simulation

| Decay rate | 1 | 0.5 | 0.3 | 0.15 |
|---|---|---|---|---|
| Sparsity level | 0.61 | 0.73 | 0.78 | 0.88 |

**Supplementary Table 3.** Gene markers used to annotate cell types in the mouse retina data set.

| Cell type | Gene markers | | |
|---|---|---|---|
| Horizontal cells | *LHX1* | *PAX6* | |
| Retinal ganglion cells | *SLC17A6* | *PAX6* | |
| Amacrine cells | *PAX6* | *GAD1* | *SLC6A9* |
| Cones | *OPN1MW* | | |
| Bipolar cells | *VSX2* | | |
| Muller glia | *PAX6* | *VSX2* | *RLBP1* |
| Astrocytes | *RLBP1* | *GFAP* | |
| Fibroblasts | *PAX6* | *SLC6A9* | *VSX2* |
| Vascular endothelium | *PECAM1* | *KCNJ8* | |
| Pericytes | *KCNJ8* | | |
| Microglia | *CX3CR1* | | |

**Supplementary Table 4.** Genes identified as differentially expressed between distal and proximal enterocytes. First, genes significantly overexpressed ($log_2$ fold change > 0.5 and rank-sum P-value < 0.05) on distal and proximal enterocytes were identified. Second, genes solely overexpressed on distal or proximal enterocytes were selected and fold-change differences between distal/proximal were calculated on these genes. Third, genes with largest fold change differences (top ten largest $log_2$ fold change difference) were displayed here. Up: genes discovered on *H. polygyrus* (day 10); Bottom: genes discovered on *Salmonella*.

*H. polygyrus* (day 10) infection

| | | E-distal | | E-proximal | |
|---|---|---|---|---|---|
| **GENE name** | Fold Change Difference | Fold Change | P-value | Fold Change | P-value |
| *Saa1* | 4.803493 | 6.742739 | 5.95E-07 | 1.939246 | 5.89E-01 |
| *Tat* | 4.595733 | 4.926558 | 2.61E-02 | 0.330824 | 9.46E-01 |
| *Pla2g4c* | 3.035611 | 1.291513 | 6.42E-01 | 4.327124 | 3.68E-03 |
| *Ifit1* | 3.024178 | 3.867634 | 2.24E-02 | 0.843456 | 1.11E-01 |
| *Isg15* | 2.75031 | 3.486814 | 9.23E-04 | 0.736504 | 6.24E-04 |
| *Sprr2a3* | 2.496977 | 2.386638 | 8.22E-04 | 4.883615 | 1.08E-09 |
| *Gsdmc2* | 2.432536 | 0.19084 | 5.16E-01 | 2.623376 | 3.76E-21 |
| *Lrg1* | 2.416572 | 5.804827 | 2.61E-02 | 3.388255 | 1.15E-01 |
| *B3gnt7* | 2.323216 | 3.662167 | 1.79E-04 | 1.338951 | 1.12E-01 |
| *Apobec3* | 2.197011 | 0.269375 | 8.11E-01 | 2.466385 | 1.37E-02 |

*Salmonella* infection

| | | E-distal | | E-proximal | |
|---|---|---|---|---|---|
| **GENE name** | Fold Change Difference | Fold Change | P-value | Fold Change | P-value |
| *Tat* | 2.953015 | 3.176097 | 2.82E-01 | 0.223082 | 9.61E-01 |
| *Gsdmc4* | 2.532489 | -3.55085 | 1.28E-04 | -1.018361 | 1.24E-02 |
| *Saa1* | 2.366475 | 4.955453 | 9.61E-05 | 2.588978 | 2.18E-01 |
| *Chst4* | 2.115494 | 2.515421 | 9.45E-02 | 0.399927 | 7.50E-01 |
| *Gsdmc2* | 1.928261 | -2.915142 | 1.09E-01 | -0.986881 | 1.95E-01 |
| *X1810065E05Rik* | 1.735235 | -1.326567 | 4.67E-03 | 0.408668 | 5.11E-01 |
| *Igtp* | 1.73344 | 4.91691 | 1.24E-14 | 3.183471 | 5.24E-29 |
| *Ifit1* | 1.716908 | 2.551523 | 2.63E-03 | 0.834615 | 1.14E-02 |
| *B3gnt7* | 1.710502 | 1.975426 | 4.90E-02 | 0.264924 | 7.76E-01 |
| *Ly6a* | 1.618224 | 2.614956 | 9.05E-02 | 0.996732 | 5.69E-01 |

**Supplementary Table 5.** Gene markers and criteria from Allen Brain Atlas (http://brain-map.org) were used to annotate three major cell groups for mouse brain data set.

| Cell type | Criteria |
|---|---|
| Glutamatergic | max(*Slc17a6*, *Slc17a7*) > max(*Gad1*, Gad2, *Olig1*, *Gja1*, *Xdh*, *Ctss*, *Myl9*, *Slc32a1*) |
| GABAergic | max(*Gad1*, *Gad2*, *Slc32a1*) > max(*Slc17a7*, *Slc17a6*, *Olig1*, *Gja1*, *Xdh*, *Ctss*, *Myl9*) |
| Non-neuronal | Other cell types |

**Supplementary Table 6.** Gene markers used to annotate cell types for mouse brain data.

| Non-neuron | | | |
|---|---|---|---|
| **Cluster number** | Cell type | Gene marker | Reference |
| **5** | Oligodendrocyte | *Pdgfra,Olig1, Olig2* | 1 |
| **25** | Oligodendrocyte Progenitor | *Olig1, Olig2* | 2 |
| **13, 29** | Endothelia | *Flt1, Kdr* | 3 |
| **19, 33** | Smooth Muscle Cells | *Abcc9, Pdgfrb* | 4 |
| **21, 27** | Astrocyte | *Gfap* | 5 |
| **22** | Microglia | *C1qb,Tgfbr1* | 3 |
| **Glutamatergic neurons** | | | |
| **Cluster number** | Cell type | Gene marker | Reference |
| **12** | Cerebellar Interneuron | *Pax3, Mki67* | 7, 8 |
| **18, 32** | Cajal-Retzius | *Trp73, Reln* | 9 |
| **28** | Claustrum Pyramidal | *Nr4a2* | 10 |
| **0, 3, 4** | Migrating Interneuron | Dlx family, *Lhx6* | 11 |
| **2, 20** | Hippocampal Granule Precursor | *Prox1, Nrp2* | 12 |
| **8** | Cortex Layer2/Layer 3 Pyramidal | *Ntf3* | 13 |
| **GABAergic neurons** | | | |
| **Cluster number** | Cell type | Gene marker | Reference |
| **1** | Sst GABAergic Neurons | *Lhx6, Maf, Sst* | 14 |
| **6** | Htr3a GABAergic Neurons | *Htr3a* | 14 |
| **10** | Thalamic Interneuron/ Cortext Layer 6 Pyramidal | *Syt6, Six3* | 15, 16 |
| **14, 17** | Hippocampal Pyramidal Precursor | *Nrp1* | 17, 18 |
| **15** | Immune Cells? | Hemoglobin family | 6 |

**Supplementary Table 7.** Gene markers used to identify mouse epithelial cell types.

| Cell type | Gene markers |
|---|---|
| Stem | *Lgr5, Ascl2, Slc12a2, Axin2, Olfm4, Gkn3* |
| Cell cycle | *Mki67, Cdk4, Mcm5, Mcm6, Pcna* |
| Enterocyte share | *Alpi, Apoa1, Apoa4, Fabp1* |
| Enterocyte proximal | *Fabp1, Lct, Cbr1, Ephx2, Gstm3, Adh6a, Creb3l3* |
| Enterocyte distal | *Mep1a, Fgf15, Clec2h, Fabp6, Mep1a, Muc3* |
| Goblet | *Muc2, Tff3, Agr2* |
| Paneth | *Lyz1, Defa17, Defa22, Defa24, Ang4* |
| EEC | *Sox4, Neurog3, Neurod2, Sct, Cck, Gcg* |
| Tuft | *Dclk1, Trpm5, Gfi1b, Il25, Lrmp, Cd24a* |

**Supplementary Table 8.** Cell-cycle related clusters were subdivided into transit amplifying early stage, transit amplifying late stage, and enterocyte progenitor based on the ratio of cell-cycle & stem cell markers to enterocyte expression.

| **Cluster index** | | 3 | 1 | 5 | 6 | 11 | 15 | 16 | 10 |
|---|---|---|---|---|---|---|---|---|---|
| Average expression (0-1 normalized UMI count) | Stem + cell cycle | 0.06338 | 0.06986 | 0.02685 | 0.04865 | 0.03809 | 0.01823 | 0.04549 | 0.03228 |
| | Enterocyte | 0.00438 | 0.00594 | 0.00366 | 0.00677 | 0.00991 | 0.00618 | 0.02017 | 0.03688 |
| Ratio | Stem/enterocyte | 14.46535 | 11.74485 | 7.33541 | 7.18305 | 3.84254 | 2.94927 | 2.25505 | 0.87513 |
| Annotation | | Transit Amplifying early | | Transit Amplifying | | Enterocyte Progenitor | | | |
